# Multi-scale immune selection and the transmission-diversity feedback of *Plasmodium falciparum* malaria

**DOI:** 10.1101/163584

**Authors:** Thomas Holding, John Joseph Valletta, Mario Recker

## Abstract

Antigenic diversity is a key factor underlying the complex epidemiology of *Plasmodium falciparum* malaria. Within-host clonal antigenic variation limits host exposure to the parasite’s antigenic repertoire, while the high degree of diversity at the population-level requires multiple exposures for hosts to acquire anti-disease immunity. This diversity is predominantly generated through mitotic and meiotic recombination between individual genes and multi-gene repertoires and is therefore expected to respond dynamically to changes in transmission and immune selection. We hypothesised that this coupling creates a positive feedback mechanism whereby infection and disease transmission promotes the generation of diversity, which itself facilitates immune evasion and hence further infection and transmission. To investigate the link between diversity and malaria prevalence in more detail we developed an individual-based model in which antigenic diversity emerges as a dynamic property from the underlying transmission processes. We show that the balance between stochastic extinction and the generation of new antigenic variants is intrinsically linked to within-host and between-host immune selection, which in turn determines the level of diversity that can be maintained in a given population. We further show that the transmission-diversity feedback can lead to temporal lags in the response to natural or intervention-induced perturbations in transmission rates. These results will add to our understanding of the epidemiological dynamics of *P. falciparum* malaria in different transmission settings and will have important implications for monitoring and assessing the effectiveness of disease control efforts.

## Introduction

Antigenic diversity is one of the key factors underlying *Plasmodium falciparum* epidemiology. Clonal antigenic variation, whereby the parasite changes expression of the immunodominant surface proteins PfEMP1 through transcriptional switches between members of the *var* gene family, enables the parasite to evade adaptive immune responses and thus to maintain long periods of infections (Borst et al., 1995; Craig and Scherf, 2001; Kirkman and Deitsch, 2012; Peters et al., 2002; Scherf et al., 2008). Furthermore, there is usually little concordance between the *var* gene repertoires of individual parasites (Barry et al., 2007; Kraemer et al., 2007; Rask et al., 2010; Tessema et al., 2015), such that any immunity resulting from one infection might not protect against subsequent infections by different parasites. As a result, hosts living in malaria endemic regions often experience a high number of malaria infections before they acquire a degree of protection that prevents them from developing symptomatic or life-threatening disease but not necessarily from infection *per se* (Langhorne et al., 2008; Reyburn et al., 2005).

Although it is becoming increasingly clear that naturally acquired immunity to malaria is orchestrated by different responses against various parasite antigens, adaptive responses against the variant surface antigens (VSA), and the highly polymorphic surface proteins PfEMP1 in particular, are thought to play a pivotal role in protecting the host from severe infection outcomes (see e.g. (Bull et al., 1998; Chan et al., 2012)), either by directly contributing to parasite clearance or by preventing parasite sequestration and its associated pathologies (David et al., 1983; Hviid, 2010; Udeinya et al., 1983). Understanding the processes underlying the generation and maintenance of antigenic diversity and how it relates to naturally acquired immunity and population-level parasite prevalence is therefore of fundamental importance. Studies have shown that *P. falciparum* population structure is spatially heterogeneous (Barry et al., 2007; Omedo et al., 2017) and exhibit different levels of diversity depending on the specific ecology of the region (Albrecht et al., 2006, 2010; Babiker et al., 1994; Paul et al., 1995). In particular the diversity of PfEMP1 and other variant antigen families likely depends to some extent on the effective parasite population size and local transmission intensity (Albrecht et al., 2006, 2010; Chen et al., 2011).

Extensive diversity among the parasite population facilitates high reinfection rates and thus parasite prevalence, simply because parasites can more readily establish an infection even in pre-exposed hosts by displaying diverse sets of antigens. On the other hand it is important to note that diversity is not a static quantity but rather the result of processes linking within-host infection and between-host transmission dynamics. That is, antigenic diversity of *var* genes and *var* gene repertoires is mainly generated by frequent intra- and intergenic recombination events, respectively (Bopp et al., 2013; Claessens et al., 2014; Conway et al., 1999; Freitas-Junior et al., 2000; Taylor et al., 2000).

Here it is assumed that low rates of mitotic recombination during asexual replication in the blood generates new *var* gene variants. These might not necessarily contribute directly to within-host immune evasion, as seen in other antigenically variable parasites, such as trypanosomes or Babesia (Barbour and Restrepo, 2000; Deitsch et al., 2009), but will still be passed on as part of the genomic *var* gene repertoire during transmission. Meiotic recombination, on the other hand, which occurs during sexual replication inside the mosquito and operates predominantly at the genome level, is more responsible for the creation of new *var* gene repertoires when mosquitoes are infected by more than one parasite genotype. The probability of this happening is itself related to population-level prevalence and diversity because (i) hosts are more likely to carry multi-clonal infections (Gatei et al., 2015; Vafa et al., 2008), and (ii) any two hosts are more likely to be infected by parasites with different antigenic repertoires (Chen et al., 2011). Put together we would thus hypothesise that parasite prevalence and diversity are governed by a non-linear feedback mechanism whereby infections and transmission events are the sources of diversity, which itself facilitates the conditions for further infections and so on.

Epidemiological studies have shown a strong non-linear relationship between malaria transmission intensity, by means of the entomological inoculation rate (EIR), and parasite prevalence in a population, which increases steeply under low to medium transmission levels but then plateaus in more intense transmission settings (Beier et al., 1999; Okello et al., 2006; Smith et al.,2005). Mathematical models trying to elucidate this relationship often focus on the slow acquisition of immunity without explicitly taking diversity into consideration. In those frameworks it is therefore simply the feedback between prevalence and transmission, whereby higher prevalence leads to higher infection rates, which generates the observed relationship under the assumption that hosts require a high number of infections to become immune (Dietz et al., 1974; Gu et al., 2003; Killeen et al., 2000; Mandal et al., 2011; Molineaux, 1985; Smith et al., 2008, 2006). One limitation of these approaches is their inability to consider the effect of temporal fluctuations in diversity on other epidemiological processes. Given the key role diversity plays in immune evasion, pathogenesis and reinfection, it seems pertinent to investigate these relationships in more detail.

Here we aimed to shed further light on how malaria prevalence, transmission and immunity are dynamically linked via parasite diversity, which itself is governed by multi-scale immune selection and feedback mechanisms. Our results demonstrate how the observed relationship between EIR and malaria prevalence emerges naturally in our system and how the transmission-diversity feedback can lead to temporal lags in the system’s response to changes in mosquito density or biting rates.

## Methods

We developed a stochastic individual-based model explicitly accounting for host and mosquito demographics, parasite diversity as well as infection and transmission events.

### Host definition

Human hosts are modelled individually and, for reasons of simplicity, are limited to two active infections. Host demographic processes are modelled by assuming a daily probability of death, given as

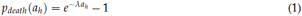

where *λ* = −1.5*e*^−6^ and *a*_*h*_ is the host age in years, i.e. we assume a constant probability of death over the course of a year. We did not account for maternal protection and assumed that upon death individuals are immediately replaced by an immunologically naive newborn to maintain a constant population.

We did not model within-host dynamics explicitly but instead calculated the length of infection as a function of the number of novel antigenic variants the parasite presents to the host (Holding and Recker, 2015), given as

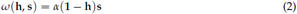

where *α* is the maximal contribution to an infection by a novel antigen (i.e. an antigen to which the host has no immunity to) and **h** = {*h*_1_, *h*_2_,…, *h*_*Amax*_} is the host’s immune status, with *h*_*a*_ representing the degree of protection against antigen type *a* (*h*_*a*_ = 1 corresponds to complete immunity to antigen *a*, and *h*_*a*_ = 0 corresponds to complete susceptibility).

In addition to strictly variant-specific immunity we also considered that exposure to variant *a* can also induce cross-protection against other, antigenically related variants, where the degree of protection decays with distance in antigen space from *a*. We implemented cross-immunity by additive transformation of the immune status vector, **h**, using a Gaussian distribution with the mean corresponding to the antigen ID and the strength of cross-immunity determined by the variance. The change in the immune status with exposure to antigen *a* can then be calculated by

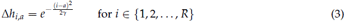

where *γ* controls the strength of cross-immunity, *R* is the size of antigen space, and *a* is the antigen the host is being exposed to. For computational simplicity the extremities of the Gaussian distribution where *h*_*i,a*_*<*0.01 were assumed to be equal to zero.

We make the simplifying assumption that upon infection the host’s immune status changes immediately to reflect exposure to all antigens in the infecting strain’s repertoire. Hence the success of future infection events is subject to these changes even if a host has not yet cleared the ongoing infection. Strains in concurrent infections are assumed to act independently with the exception of this indirect immune interference.

### Recombination

We considered diversity generation through recombination at both the gene and genome level. For sexual recombination we assumed that two strains, picked up by the mosquito from a multiclonal infection, give rise to a single recombinant strain, whose repertoire is generated from the parental repertoires in a gene-wise manner by probability *p*_*s*_. That is, we assumed that a gene can be taken from either parent strain in an independent manner. Although this might not be the most biologically realistic assumption and ignores any intra-genomic structuring (e.g. by means of chromosomal location and upstream promoters etc.), it maximises the generation of new repertoires and more easily disrupts the emergence of strain structuring.

Mitotic recombination between individual genes is assumed to occur during asexual reproduction in the host (Bopp et al., 2013; Claessens et al., 2014). It is expected that parasites carrying a novel gene resulting from mitotic recombination only make up a small fraction of parasites in the host, and that the magnitude of change is so small that there would be little bearing on an ongoing infection. Additionally, we expect the probability of mosquitoes picking up these parasites to be much lower than the more numerous original clone. For computational reasons we do not compute this at the within-host level, but instead assumed a small probability, *p*_*c*_, that a recombinant gene is copied to the transmitted strain. *p*_*c*_ therefore incorporates the probability that a recombinant parasite is taken up during the blood meal as well as the rate of intragenic (mitotic) recombination.

Mitotic recombination is implemented by adding a number, *r*, to the original antigen representation, where *r* is proportional to the difference between the original and donor antigens, given as

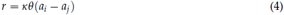

where *κ* is a random number drawn from a uniform distribution *U* (−1, 1), *θ* scales the magni-tude of change, and *a*_*i*_ and *a*_*j*_ are the original and donor antigen types, respectively. The new recombinant gene is then represented as

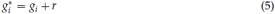

where *g*_*i*_ is the full genotype representation (not antigen type) of the parent antigen.

This scheme avoids having to simulate the low-level biological mechanisms, such as insertion and deletion of domains or nucleotide sequences between genes, leading to a computationally efficient model which emulates the main features of recombination: (i) recombination between antigenically similar donor genes is likely to produce recombinant genes which are similar to the donor genes, (ii) recombination between antigenically dissimilar genes more likely results in a recombinant antigen which differs from the original genes, and (iii) recombination events alter the genotype but do not necessarily change the antigenic type, although these silent changes can accumulate over time and eventually lead to changes in the antigen type.

### Mosquito definition and transmission

Mosquitoes are modelled individually and can be uninfected, exposed or infected and infectious with a single parasite strain. Mosquitoes are only assumed to die of natural causes and are immediately replaced with new uninfected individuals to maintain a constant population. The age-related probability of death is modelled using a logistic function

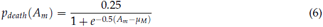

where *μ*_*M*_ is the mean lifespan and *A*_*m*_ is the mosquito’s age in days.

Mosquitoes are assumed to bite at a constant rate, *b*. When an infectious mosquito bites a host it transmits the infection unless the host is at its maximum capacity for concurrent infections. Equally, an uninfected mosquito which feeds on an infected host becomes infected with a fixed probability, *p*_*trans*_. Unlike humans, mosquitoes can only be infected by a single strain. For simplicity we ignored the human incubation period but considered it for the mosquito, due its much shorter lifespan. Infected mosquitoes are assumed to stay infected for life.

### Initialisation

The model was initialised with both human and mosquito populations at a demographic equilibrium. A small number of mosquitoes were initialised infected while all humans were initialised as naive (i.e. with no prior exposure). As a result, a burn-in period is required to allow the system to reach a dynamic equilibrium. We generated an initial pool of antigens of size *A*_*init*_ from a uniform distribution over the entire genotype space. Next we generated *S*_*init*_ strains by randomly sampling from the antigen pool without replacement (or with replacement in cases where *A*_*init*_*<S*_*init*_), which were then used to infect mosquitoes, such that an approximately equal number of mosquitoes were infected by each initial strain.

Table 1 summarises the main parameters used in our model and their baseline values

**Table 1:**
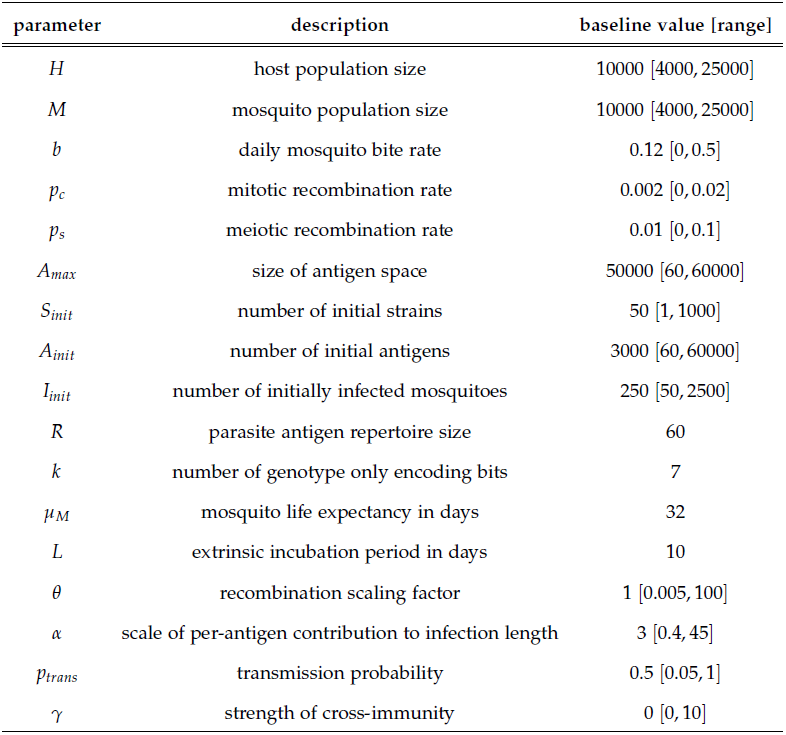
Main model parameters.

| parameter | description | baseline value [range] |
| --- | --- | --- |
| $H$ | host population size | 10000 [4000, 25000] |
| $M$ | mosquito population size | 10000 [4000, 25000] |
| $b$ | daily mosquito bite rate | 0.12 [0, 0.5] |
| $p_c$ | mitotic recombination rate | 0.002 [0, 0.02] |
| $p_s$ | meiotic recombination rate | 0.01 [0, 0.1] |
| $A_{max}$ | size of antigen space | 50000 [60, 60000] |
| $S_{init}$ | number of initial strains | 50 [1, 1000] |
| $A_{init}$ | number of initial antigens | 3000 [60, 60000] |
| $I_{init}$ | number of initially infected mosquitoes | 250 [50, 2500] |
| $R$ | parasite antigen repertoire size | 60 |
| $k$ | number of genotype only encoding bits | 7 |
| $\mu_M$ | mosquito life expectancy in days | 32 |
| $L$ | extrinsic incubation period in days | 10 |
| $\theta$ | recombination scaling factor | 1 [0.005, 100] |
| $\alpha$ | scale of per-antigen contribution to infection length | 3 [0.4, 45] |
| $p_{trans}$ | transmission probability | 0.5 [0.05, 1] |
| $\gamma$ | strength of cross-immunity | 0 [0, 10] |

## Results

We investigated the effect of antigenic diversity on malaria transmission and prevalence by means of a stochastic individual-based model (see Methods) in which antigen variants are dynamically generated through recombination and in which the diversity of the parasite population emerges from the underlying transmission dynamics. In order to highlight how population-level prevalence and transmission rates are related to parasite diversity we first show the model behaviour assuming semi-static levels of antigenic diversity, i.e. without accounting for recombination, before investigating the dynamical feedback between parasite antigenic diversity and malaria epidemiology.

### Static diversity

Strains were initialised by randomly selecting antigens without replacement to ensure that there is no overlap between repertoires. This allowed us to assess the general effect of diversity without the added complications of immune interactions. Without the possibility of new variants entering the population, the model converges towards a semi-equilibrium dictated by the background transmission rate, here measured as the daily biting rate, and the total level of diversity among the parasite population (figure 1a).

**Figure 1:**
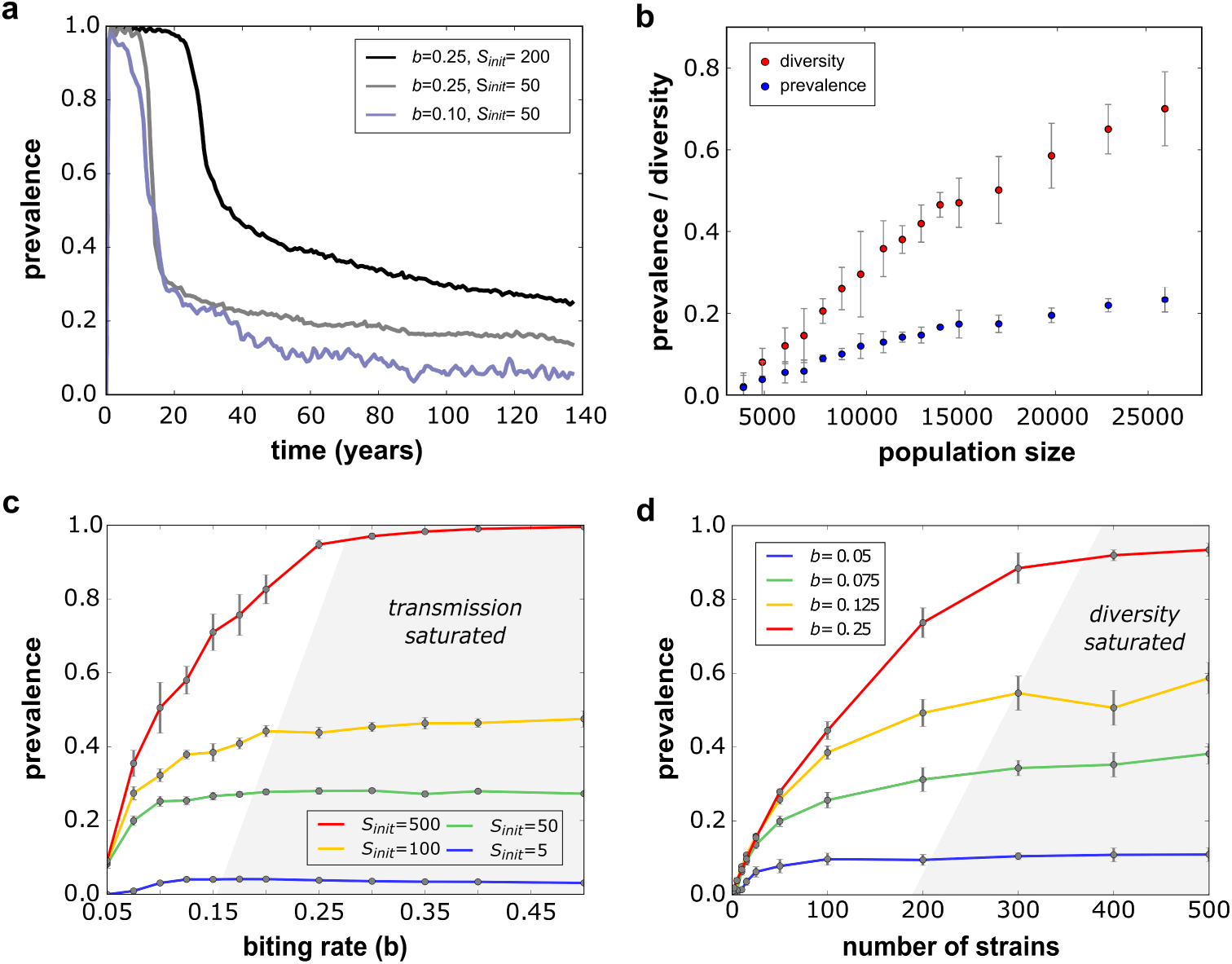
Relationship between transmission potential, antigenic diversity and malaria prevalence. (**a**) Simulated timeseries showing how malaria prevalence, defined as the proportion of the population infected by the parasite, converges towards an endemic equilibrium determined by the daily biting rate and antigenic diversity. (**b**) Diversity, here measured as the proportion of initially circulating antigenic variants that are maintained in a population, is positively correlated with the size of the host population (assuming equal *M*:*H* ratios), which also affects the equilibrium levels of malaria prevalence. (**c**) Equilibrium levels of malaria prevalence as a function of the transmission potential (biting rate) under different levels of antigenic diversity. In all cases, prevalence plateaus and does not increase further with increasing biting rates; we refer to this regime as *transmission saturated*. (**d**) Equilibrium levels of malaria prevalence as a function of diversity under different levels of transmission, showing a plateauing behaviour where prevalence does not increase any further with increasing levels of diversity; we refer to this regime as *diversity saturated*. Results for (**b**)-(**d**) based on 10 model runs, with error bars indicating the standard errors around the mean. Parameter values, unless stated otherwise: *H* = 10000, *M* = 10000, *b* = 0.12, *S*_*init*_ = 50, *A*_*init*_ = 3000.

A crucial factor influencing the relationship between malaria prevalence and parasite diversity is the size of the host population. As in this model formulation we did not account for the generation of diversity over the course of our simulations, individual variants and parasite strains were subject to stochastic extinction. In fact, in each model run we observed that only a certain proportion of the initial set of variants were maintained over a given period of time. Larger populations are known to be able to maintain higher degrees of diversity and we see the same phenomenon in our model. That is, we find a strong positive correlation between host population size and the proportion of initial antigens retained and hence an overall but small increase in prevalence (figure 1b).

As expected, for a given number of antigenic variants that co-circulate in the population there is a positive but non-linear relationship between mosquito biting rate and population-level parasite prevalence, here defined as the proportion of the population that is currently infected. After an initial steep increase in prevalence with increasing rates of transmission the relationship plateaus, up to a point where further increases in transmission does not raise parasite prevalence any further (figure 1c). This scenario, which we refer to as *transmission saturated*, occurs as hosts acquire immunity to the vast majority of the antigenic variants available in the population. Therefore, the attained equilibrium rate in prevalence is strongly dependent on antigenic diversity, with higher levels of diversity enabling the parasites to more readily find susceptible hosts, leading to a higher proportion of infected individuals (figure 1c).

A similar relationship can also be found between antigenic diversity and parasite prevalence with increasing levels of diversity leading to an increase in prevalence, at least up to a point where further diversification does not affect prevalence any more. This scenario, here referred to as *diversity saturation*, occurs when the number of malaria infections a host acquires over their life-time is simply limited by exposure. As a result, and pretty much as expected, higher levels of exposure, i.e. biting rates, will shift the equilibrium levels of parasite prevalence upwards, as shown in figure 1d.

### Dynamic diversity

As clearly demonstrated in figure 1, there is a strong link between parasite antigenic diversity, i.e. the degree by which the parasite can circumvent immune responses and establish infections even in pre-exposed and semi-immune individuals, and malaria prevalence. In the examples shown above it was assumed that diversity was an initially fixed but slowly declining quantity. In reality, however, antigenic diversity in *P. falciparum* malaria is the result of dynamic processes, namely recombination and mutation, whose rates are determined by epidemiological parameters related to transmission and prevalence.

To demonstrate the effect of considering diversity as a dynamic property we ran our model without recombination for a number of years until it reached a semi-equilibrium state before turning recombination on. As clearly illustrated in figure 2, allowing new variants and parasite strains to be generated over time leads to a significant increase in overall diversity, here defined as the percent of all possible antigenic variants, *A*_*max*_ (figure 2a). This increase in diversity effectively reduces population-level immunity, as hosts will not have experienced the newly generated variants before (figure 2b). As a result, parasites will find it easier to find susceptible hosts, leading to an increase in parasite prevalence rates (figure 2c) and hence disease transmission (EIR, figure 2d), even without changes to the transmission potential. Not surprising, we found a strong correlation between the recombination rate and the system’s response with regards to these epidemiological determinants.

**Figure 2:**
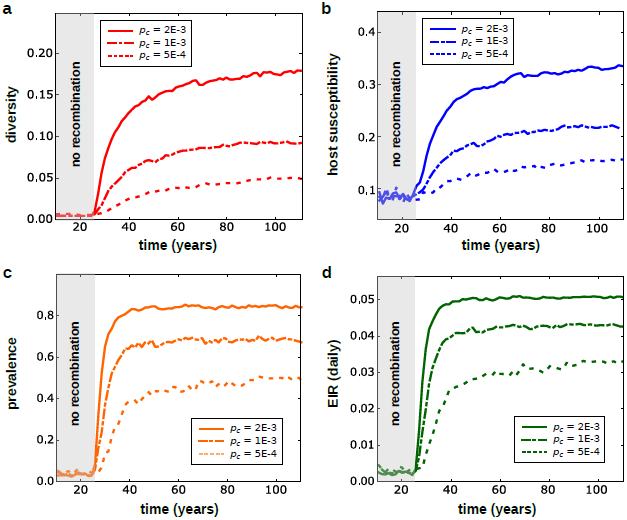
Diversity as an emergent property of infection and transmission. (**a**) Allowing for recombination to create new antigenic variants and antigenic repertoires significantly increases the level of diversity amongst the parasite population, here defined as the % of the assumed maximum level of diversity. (**b**) As diversity increases, host susceptibility increases as parasites carrying novel variants find it easier to re-infect individuals with prior immunity. (**c**) Increasing diversity and hosts susceptibility leads to higher malaria incidence and population-level prevalence. (**d**) Increasing the number of infected hosts increases the overall transmission intensity (EIR) even without changes to the biting rate. Different lines denote different rates of recombination (*ρ*), showing how higher rates of diversity generation relate positively with parasite prevalence and disease transmission. Parameters values: *M* = 10000, *H* = 10000, *A*_*max*_ = 50000, *A*_*init*_ = 3000, *S*_*init*_ = 50, *b* = 0.12, *p*_*s*_ = 0.01, *γ* = 0

Figure 2 clearly demonstrates the positive feedback between parasite antigenic diversity and disease prevalence in the population. What is also apparent from these graphs is that this process is not unbounded but that the system will settle onto a new equilibrium that is balanced between diversity generation, determined by the rates of mitotic and meiotic recombination as well as background transmission rates, and diversity loss, due to demographic and immune selection associated risk of extinction. Immune selection in particular has a strong and expected effect on both antigenic diversity and parasite prevalence. That is, theoretical models have repeatedly shown how cross-immunity can structure antigenically variable pathogen populations into sets of strains with non-overlapping antigenic repertoires (Gupta and Anderson, 1999; Gupta et al., 1996). In our model, increasing the degree of cross-immunity that each variant antigen elicits against antigenically similar variants enhances selection pressure on the pathogen to find susceptible hosts, leading to an increased risk of extinction and thus a decrease in overall diversity and parasite prevalence. This is demonstrated in figure 3, where we simulated our model using the same assumption about transmission and recombination under increasing immune-selection pressure (degree of cross-immunity, *γ*).

**Figure 3:**
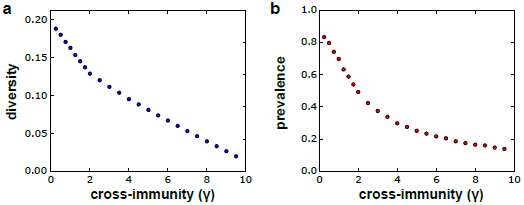
Antigenic diversity and malaria prevalence as a function of immune selection pressure. Cross-immunity determines the degree of inhibition that each antigenic variant elicits against antigenically similar variants, such that higher levels of cross-immunity increases the selection pressure on the parasite population, which in turn limits the number of variants that can be maintained in a population (**a**) and thus decreases the overall level of malaria prevalence (**b**). Each point is the average equilibrium level based on 10 model runs. Parameters values: *M* = 10000, *H* = 10000, *A*_*max*_ = 50000, *A*_*init*_ = 3000, *S*_*init*_ = 50, *b* = 0.12, *p*_*c*_ = 0.002, *p*_*s*_ = 0.01

### Diversity and prevalence under changing transmission rates

As diversity and prevalence are coupled dynamically and are partially determined by the transmission potential in terms of mosquito biting rate, we hypothesised that there must be a lag in the system’s response to temporal changes in disease transmission, caused for example by changes in mosquito population density or changes in bednet usage. We analysed this by increasing or decreasing the biting rate over a period of four years and recorded the resulting response in antigenic diversity (figure 4a and c) and parasite prevalence (figure 4b and d) over time.

**Figure 4:**
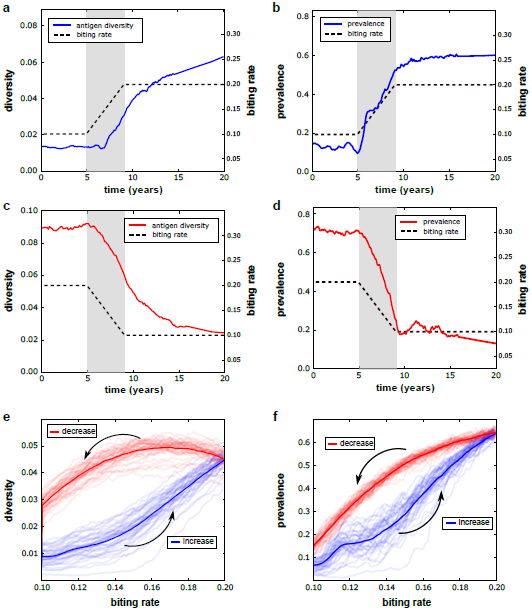
Antigenic diversity and malaria prevalence in response to changes in transmission. As antigenic diversity and parasite prevalence are linked via a dynamic feedback loop, our model predicts a temporal lag in the response to both increases (**a** and **b**) and decreases (**c** and **d**) in transmission rates (mosquito biting rate, black dashed lines). The periods over which the biting rate was changed is highlighted in grey. The system also exhibits a degree of inertia, with changes in diversity and prevalence taking place many years after the biting rate has settle onto a new value. The different rates at which the system responds to changes in the transmission rate can results in different levels of diversity (**e**) and prevalence (**f**), depending on whether there has been a reduction (red lines) or increase (blue lines) in transmission, at least when monitored over the period where the change is taking place. Results are shown for 100 model runs, with the bold lines showing the average levels. Parameter values: *M* = 8000, *H* = 8000, *p*_*c*_ = 0.001, *p*_*s*_ = 0.01, *A*_*init*_ = 2400, *S*_*init*_ = 40, *A*_*max*_ = 50000.

In both cases of increasing and decreasing transmission potential the model showed a predictable response, with higher transmission rates leading to higher levels of diversity and prevalence rates and *vice versa*. However, and in particular in those cases where we simulated an increase in transmission (figure 4a and b), we also observed a certain inertia, where both diversity and prevalence kept increasing for many years despite no further changes to the biting rate. This can be explained by the positive feedback loop between diversity and prevalence, where a change in one property has delayed downstream effect. Interestingly, though, we found that the system would generally respond quicker to decreases in transmission, although even in those cases it took many years for the system to attain a new state of equilibrium.

The system’s intrinsic inertia also leads to the phenomenon of hysteresis, where different rates of transmission can have very different outcomes in terms of diversity and prevalence depending on whether there has been an increase or a decrease in the biting rate. This is shown in figures 4e and f, which plot the levels of diversity and parasite prevalence during the transition periods from low-to-high (blue lines) and high-to-low (red lines) mosquito biting rates. What is clear from these graphs, and figure 4 in general, is that the relationship between malaria prevalence and other external factors that could influence its transmission potential is highly non-linear and time-lagged to the point where observed changes in malaria incidence, for example, could be due to changes in mosquito abundance that had happened a considerable period of time back in the past.

## Discussion

Here we analysed the diversity-transmission feedback and its implication for the epidemiological dynamics of *P. falciparum* malaria. Our model explicitly considered antigenic diversity as a dynamic property, balanced between the continuous generation of new variants through recombination and stochastic extinction. Malaria transmission models have a long history going back to the original Ross and Macdonald framework (Macdonald et al., 1950; Macdonald, 1952; Ross, 1905), which has formed the backbone of many theoretical attempts to understand the relationship between the parasite, its host and the mosquito vector (reviewed in Smith *et al.* (Smith et al., 2012)). Based on the apparent absence of complete immunity against malaria in the field, many frameworks either do not consider acquired immunity at all (as in the original model formulation) (Killeen et al., 2000; Macdonald et al., 1950; Macdonald, 1952; Ross, 1905; Smith et al., 2006) or implicitly include it as an exposure-dependent reduction in the risk, duration and / or transmissibility of an infection (Dietz et al., 1974; Gu et al., 2003; Smith et al., 2008).

Naturally acquired immunity to *P. falciparum* malaria is a complex and poorly understood process. It is generally recognised that sterile immunity, which effectively prevents an infection, may never be acquired but that hosts living in malaria endemic regions build up a degree of clinical immunity that protects them first from life-threatening illness and eventually from symptomatic disease in an exposure-dependent manner (Gupta et al., 1999; Hviid, 2005; Langhorne et al., 2008). What is currently not known is what effect the reduced risk of clinical malaria has on the transmissibility of subsequent infections. That is, although there is an overall trend towards lower levels of parasitaemia and sexual parasite densities in semi-immune individuals, it is clear that even chronic, asymptomatic infections in older hosts are sufficiently transmissible (Bousema et al., 2014; Laishram et al., 2012; Lindblade et al., 2013; Okell et al., 2012).

Here we considered an alternative formulation of immunity that explicitly depends on the host’s exposure to the parasite’s variant surface antigens (VSA). Importantly, this formulation relaxes previous assumptions about the number of infections required for a host to acquire immunity. In fact, in our model this arises naturally through the interplay of antigenic diversity and transmission intensity. The role of VSAs, and PfEMP1 in particular, in both malaria pathology (Kaestli et al., 2006; Kirchgatter and Portillo, 2002; Kraemer and Smith, 2006; Kyriacou et al., 2006; Rottmann et al., 2006; Salanti et al., 2004) and acquired immunity (Bull and Abdi, 2016; Bull and Marsh, 2002; Warimwe et al., 2009) is well documented, and their extensive sequence and antigenic diversity can be seen as one of the reason why anti-disease immunity is such a slow process that relies on repeated and probably continuous exposure to the malaria parasite. We made the simplifying assumption that immunity is solely dependent on exposure to VSA, thereby ignoring the role of many of the other malaria antigens or other immunological processes that might contribute to acquired protection. However, as the purpose of this work was to highlight and qualitatively investigate the effect of antigenic diversity and how it is dynamically related to disease transmission and population-level prevalence, the contribution of other immune factors is not expected to change the results presented here, at least not in a qualitative way.

The crucial point of our model was to consider antigenic diversity not as a static quantity but rather as a dynamic property of the system that emerges naturally through multi-scale processes related to infection, transmission and immunity. That is, we considered these processes linked by a positive feedback loop. Under this assumption, diversity, in terms of novel antigenic variants, can be generated through mitotic recombination during bloodstage malaria infection. These new variants, if transmitted, are disseminated throughout the parasite population by means of meiotic recombination, which we assumed acts only at the repertoire level. This in turn will help the parasite to circumvent pre-existing immunity and thus facilitate the generation of further diversity. Importantly, the continuous generation of new antigenic variants does not result in ever-increasing levels of diversity but is counter-balanced by the loss of diversity due to stochastic extinction, where small host populations and strong immune selection pressure significantly increase the risk of parasite strains going extinct. Assuming that mitotic recombination rates and immune interference between antigenic variants are intrinsic properties of the parasite and the host, respectively, our results thus suggest that each system, potentially representing different malaria endemic settings, will attain their own state of equilibrium with regards to parasite diversity and population-level prevalence, with host and vector population sizes being the most influential factors.

The feedback between infection and diversity not only leads to temporal lags in the responsiveness of the system to changes in the transmission potential but also introduces a certain degree of inertia. Delays in a dynamical system’s response to external perturbations are expected; however, in our case we observed transient behaviours in parasite prevalence and diversity many years after the assumed changes in mosquito biting rates. In epidemiological terms this implies that trying to infer the causative factors of observed changes in malaria incidence might be more complicated than previously appreciated and would have to take into account potential changes that took place many years in the past. Together with the possibility that in a given endemic setting the system could be in a state of *transmission saturation* (figure 1b), evaluating the effectiveness of control measures directed against the mosquito (i.e. mosquito control or bednet usage) could potentially show up significant discrepancies, at least in cases where the evaluation period is too short for the system to reach a new endemic equilibrium.

There are various caveats in our model, mostly related to our simplifying assumptions about how hosts acquire immunity and how antigenic variants contribute to infection length and thus a strain’s probability of onward transmission. At the moment, very little is known about how acquired immunity relates to transmission success, although a general decrease with repeated exposure would be a reasonable assumption. In this respect, shortening the infectious period and keeping the per bite transmission probability constant is akin to lowering the transmission probability for the same length of infection. Furthermore, adding a non-VSA associated immunity component, where the risk or length of infection simply decreases as a function of accumulated exposure is not expected to lead to significantly different outcomes, thus making our results fairly robust to changes in the underlying assumptions about immunity.

## Conclusion

In summary, we have shown that antigenic diversity plays a crucial role in the epidemiology of *P. falciparum* malaria. By allowing diversity to be an emergent property of the dynamic processes underlying infection and transmission events we have demonstrated how population-level parasite prevalence and incidence are specific to an epidemiological setting and crucially determined by host and vector demographies. This would argue against a *one-model-fits-all* approach to modelling malaria epidemiology and control and also makes the case for a concentrated effort to further our understanding of the relationship between antigenic diversity, naturally acquired immunity and parasite transmission.

